# The Three Stages of Learning to Master a Sensory Augmentation Device: Activation - Acquisition - Integration

**DOI:** 10.1101/2024.07.17.603884

**Authors:** Sascha Mühlinghaus, Vincent Schmidt, Peter König

## Abstract

By augmenting sensory perception through technology, researchers study how humans use and perceive the novel sensory input provided. Nevertheless, little is known about how learning unfolds over time. To this end, 27 participants trained with an augmentation device (feelSpace belt), which provides tactile feedback about the direction of the cardinal north, over the course of six weeks. During the training phase, we tracked participants’ progress using two different self-report questionnaires containing Likert-scale and open questions. Experts quantified responses to open questions using a previously established category system. As human raters are known to be susceptible to biases, we later reproduced the expert categorization using ChatGPT-4, finding a high congruence between the two classification approaches. The results suggest a three-stage model that best describes the process of acquiring an augmented sense. During the early activation stage, processing the augmented signal requires effort, induces fatigue, and a heightened awareness of the environment. In the knowledge acquisition phase, participants develop a more detailed cognitive spatial representation containing information about objects and places in relation to their current location. The deep integration stage is marked by participants seamlessly integrating the augmented information into existing perceptual processes and automatic use. The results support the supervised use of large language models (LLMs) for the analysis of qualitative data.

## Introduction

Understanding how humans learn to use the sensory feed-back they process to predict and select appropriate actions is crucial to understanding perception and how the human brain interacts with the body and the environment. By interacting with the environment and processing sensory feedback, one may form a representation that enables purposeful actions. O’Regan and Noë (1) posited that perception occurs through the mastery of sensorimotor contingencies (SMCs), i.e., the knowledge of how sensory information changes when acting in the world. They further assert that the relationship between action and sensory feedback is reciprocal and consistent. For instance, the same action should produce the same perception under identical conditions. Therefore, agents can acquire such SMCs and perceive them as law-like relationships. Ultimately, mastering SMCs enables them to predict their environment by relating changes in their perception while accounting for their physical actions. This process may be subconscious. To study SMCs, it is relevant to consider that senses are usually acquired from a young age. Thus, it is a challenge to gather valuable insights into the perception of humans while they are still actively obtaining a new sense. Studying the acquisition of a novel sense has been suggested (2, 3) as a promising solution to gain a deeper understanding of the emergence of a reciprocal, rule-like relationship between action and sensory feedback.

Approaches to extend or enhance human perceptual abilities through technology, such as creating a novel sense, are termed sensory augmentation. As opposed to sensory substitution, where relevant information from a missing sense is provided through a different sensory modality, augmentation refers to enhancing an existing or creating a novel sense (4). Examples include augmented and virtual reality (VR) in the visual domain (5, 6), the manipulation of the tactile domain (7), and also tactile augmentation of spatial cues (2, 3, 8) or the mapping of sounds to real-time data recorded by external sensors in the auditory domain (9, 10). These successful examples of sensory augmentation show that, when implemented in a meaningful, comprehensive way and given participants train sufficiently, novel senses can be learned and integrated with existing senses in an action-related context. The specific context and what kind of augmented sensory information is processed and learned is also important to consider when studying the acquisition of a novel sense, as it would likely affect the creation of SMCs. Previous research showed qualitative and quantitative differences in how well individuals can process information from the augmented modality (11, 12). A fairly direct application to study the effect of obtaining a novel sense is the field of space perception. By moving through the environment, humans constantly perceive changes as sensory feedback, which, in turn, directly or indirectly affect their subsequent actions and perception. The “EyeCane” is a device that enables blind or visually impaired (BVI) users to hear the distance of objects in space through auditory augmentation of a distance signal recorded by a sensor (13). “EyeMusic” (9), on the other hand, aims to be applied in a wider context, allowing for more detailed object recognition. Compared to a single signal, such as distance, various signals related to objects’ size, color, and shapes within the visual scene require simultaneous translation. The feelSpace belt is a tactile augmentation device that functions like a tactile compass by providing constant feedback regarding the user’s orientation towards the cardinal north. The first study using this device was published by Nagel and colleagues (2). It investigated how the augmented signal was integrated, processed, and perceived. The authors concluded that spatial cognition and perception can be affected profoundly (2). Kaspar and colleagues (3) introduced an additional weekly questionnaire. The results show that sighted participants reported altered space perception and improved navigation abilities after seven weeks of daily wearing and training, as opposed to a control group. Brain imaging data additionally supports the results. First, sleep-electroencephalography (EEG) data showed increased brain activity related to procedural learning during the first weeks of training (14). Second, functional magnetic resonance imaging (fMRI) during a ‘path integration’ task revealed different activations in sensorimotor and higher-level motor brain areas. Schmidt and colleagues (11) used a custom-made VR environment, a larger sample size, and a large variety of spatial tasks performed directly within the VR environment. The results showed a significant increase in route and survey knowledge after six weeks of training for the belt group as opposed to the control group. Overall, sensory augmentation has been shown to lead to changes in brain activity, use of spatial strategies, improved performance on spatial tasks, and changes in the perception of space.

While previous research has offered initial insights into the mechanisms behind successful SMC formation, the underlying temporal dynamics are uncertain. As the presented findings only focus on the difference before and after training, the exact temporal dynamics of this learning process can often not be investigated further. Moreover, studies usually involve specific contexts that suit the intended use case of the augmentation device. Therefore, it is difficult to generalize results on learning times to other fields of application. Gaining a better understanding of variations in learning rates and their external causes will also benefit the study of interindividual differences in human sensory perception. Perception, as opposed to action, cannot be measured directly since it involves qualia (15). Approaches to measuring perception, therefore, usually involve indirect self-report measures such as questionnaires. While these have proven useful tools, they can be prone to human bias or manipulation (16, 17). Therefore, authors of a previous study (3) developed a category system that would allow for participants’ written answers to open questions to be quantified by two independent expert raters so that inter-rater reliability could be computed. While this approach can help gain insights into perception, it would require more resources like raters and participants to validate the resulting category system. Sensory augmentation has been shown to lead to a different space perception using various self-report measures useful to gain novel insights, yet often hard to validate and potentially unreliable.

The issue of subjective bias and error in human-rater-based categorization could be addressed using an automated classification. A more neutral approach to generating and classifying language may be presented within the domain of large language models (LLMs) designed to find statistical patterns in text documents (18). The general applicability of GPT 4 (19) poses a chance for efficient classification according to a given scheme. Moreover, it allows sharing prompts or even entire chats with the public, which could contribute to more transparent and reproducible research. To verify the findings on the changes occurring during sensory augmentation, LLM classification abilities indicate a method for neutral classification of the qualitative data.

Here we investigate perceptual and navigational changes at close intervals throughout the training period related to wearing the feelSpace sensory augmentation device. We address the difficulty of objectively assessing perceptual changes with physical measurement techniques by focusing on selfreported data classified to obtain quantified measures. This approach is verified by a classification involving ChatGPT 4. We aim to describe the temporal dynamics and corresponding perceptual changes underlying the acquisition of a novel sense.

## Methods

In this study, we collect and analyze data on perceptual changes during sensory augmentation to better understand the learning process when using augmented senses. Specifically, we evaluated the responses of participants who were learning to use a belt that indicates cardinal north through tactile stimulation around the waist. Responses were recorded during six weeks, which we will refer to as the training period. We monitored the participants’ learning process with daily and weekly questionnaires (see Data Availability). After the training period, while the participants continued to use the belt, an elaborate behavioral experiment followed in which we assessed the participants’ spatial knowledge acquisition compared with a control group. The results of these measurements after the training period are published in Schmidt et al. (11). Here, we will focus on the previously unpublished data recorded during the training period. We aim to provide more insights into the perceptual and cognitive processes underlying the acquisition of a novel sense during active training, and the underlying temporal dynamics of this learning process. During the training period with the sensory augmentation device, the participants were closely monitored to register changes regarding space perception and navigation.

### Participants

Participants were recruited via mailing lists, flyers, and suitable social media platforms, as described in Schmidt et al. (11). Informed consent was obtained from all participants after they had received a general introduction to the study. We handled the selection of participants based on a general screening questionnaire. Exclusion criteria involved age above 39, current or past substance abuse or addiction, medical abnormalities that could impact cognitive functions, use of psychotropic drugs, and history of neurological conditions or head injury. One participant dropped out during the training period for undisclosed reasons. The final sample contained 27 healthy adults. The sample was drawn from the city of Osnabrück and the surrounding area. These individuals served as participants for a monetary reward or university-internal participation credits. All participants were fluent in English or German. All study materials were provided in English or German given the participant’s preference. The ethical committee of Osnabrück University approved the study’s protocol before the recruitment.

### Materials

Sensory augmentation was achieved using the feelSpace belt (2, 3, 11). The feelSpace belt is a vibrotactile sensory augmentation device in the form of a belt worn around the waist. To provide augmentation, sixteen Vibro motors are evenly spaced around the user’s waist. It contains an integrated compass and control unit. While the controller allows various modes, we set it up so the belt serves as a tactile compass. Only one vibration motor facing the direction of the cardinal north was activated at a time. The feelSpace belt provided a real-time augmented signal of a person’s orientation towards the cardinal north through vibration around the waist.

To get used to the augmented signal, all test subjects participated in a six-week training phase where we tracked their progress with two questionnaires. The aim was to measure training progress and perceived changes. Both questionnaires were adapted from Kaspar and colleagues (3). One was assessed daily, and the other one once per week (see Data Availability). For instance, participants were asked to track their daily activities with the belt, their active training time, and their daily belt-wearing duration. We used the responses to these self-report questionnaires to calculate participants’ daily belt-wearing duration and active training time (i.e., running or cycling) with the augmented sense. Moreover, the questionnaires included 5-point Likert scale items monitoring, for instance, participants’ motivation to wear the augmented sense. Finally, the questionnaires included open questions, for example, regarding potential changes participants observed in their perception of space or the signal of the augmented sense. All responses will be analyzed as described in the Data Analysis section, to gather further insights into the temporal dynamics of acquiring a novel sense.

### Procedure

The experimental setup in this investigation is the belt training, participants underwent a 6-week training period wearing the feelSpace belt. To learn to apply the belt’s signal, participants must engage in sufficient physical movement. We asked the participants to wear the belt as much as possible throughout the day. Additionally, there were daily obligatory physical activities, such as hiking, running, and cycling, set to be at least 1.5 hours. Questionnaires surveyed the progress throughout the training period, monitoring the participants’ training progress and any potential issues (see Data Availability). Our team also called them daily to check their status. Moreover, each evening the participants filled out an online questionnaire where we asked them to report and reflect on their activities, perceptions, and feelings throughout the day. Lastly, the training progress was assessed at the end of each training week by filling out extensive online questionnaires, which were filled in with an experimenter present to discuss potential technical issues. These measures secured fully functional operations with detailed monitoring during the training period.

### Data Analysis

As various types of data were collected, we applied appropriate analytical approaches for each type. For Likert scale data, which the Shapiro-Wilk test indicated was not normally distributed, we utilized the Wilcoxon signedrank test. This non-parametric test assesses differences between two measurements in a within-subjects design. Test statistics and *p*-values for all tests conducted on the numerical data are detailed in the Results section (also see Data Availability). Note that the reported values have not been adjusted for multiple comparisons.

To obtain a quantitative measure of the responses to open questions in the weekly questionnaire, we adopted a category system that has been suggested by previous research (3, 20). The category system aims to classify reports on perceptual changes during sensory augmentation (see Fig. 1). It consists of three levels: the main-category level (5 categories), the first-order (18 categories), and the second-order category levels (56 categories). The main level comprises the categories “Space perception”, “Belt perception/experience”, “Navigation” and “Feelings”. On the main-category level and within each category of the first-order level, there is also a “Residuals” for statements that cannot be assigned to one of the given content categories. The first-order categories are entailed in the main-category level and the second-order categories are entailed in the first-order categories. This structure enables a progressively more specific classification. Statements assigned to a main category and a first-order category that cannot be classified further into a second-order category are instead assigned to the “Residuals” of the respective firstorder category. Guided by the category system, two independent raters categorized the statements with the MAXQDA software. Once each rater had independently assigned categories to the reports of a participant, the raters compared their classification. In case of incongruencies, the statement was discussed and an agreement was found to align the classification. By categorizing the data, we can compile frequency counts for each category, enabling us to draw quantitative inferences from qualitative information.

**Fig. 1.**
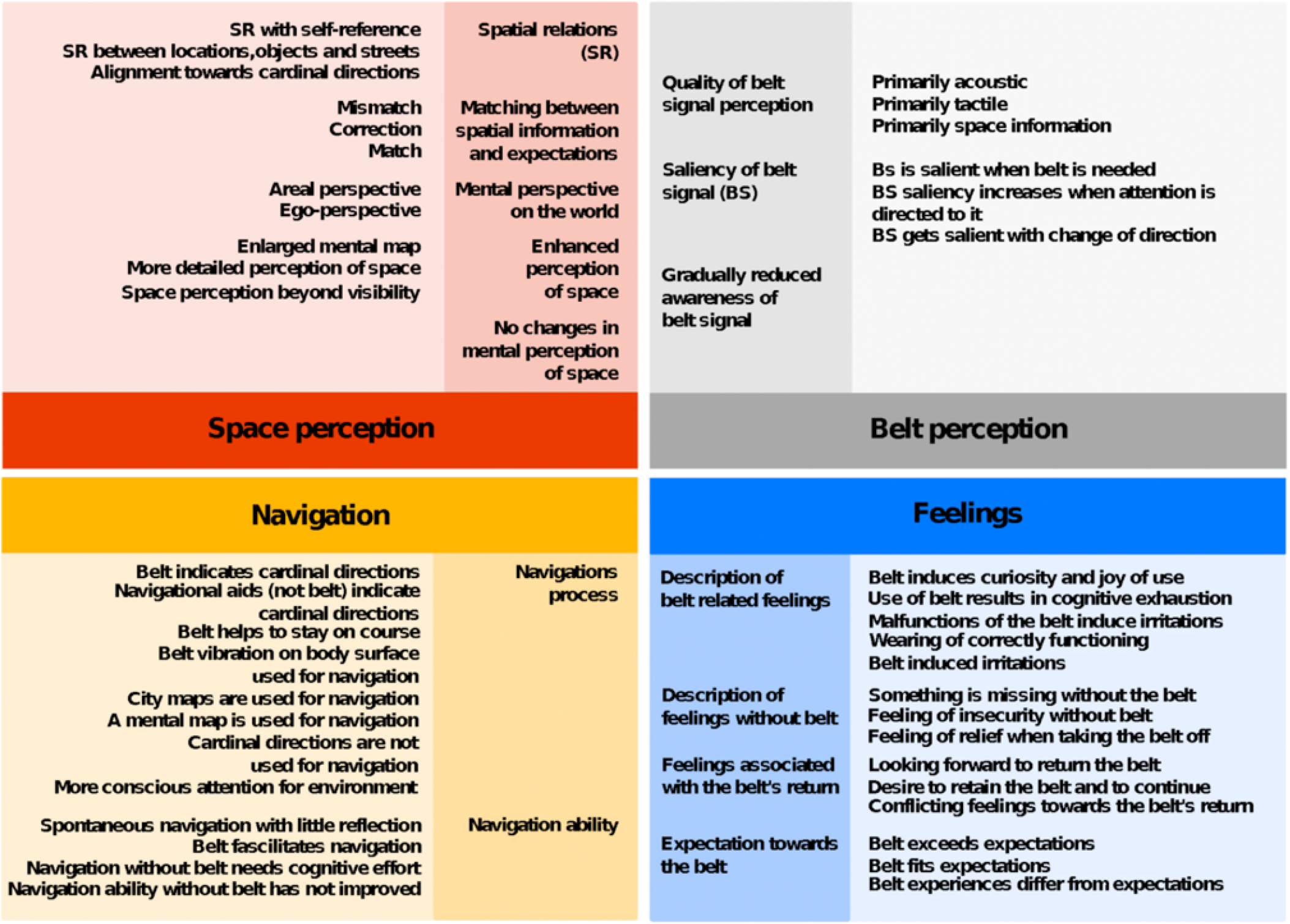
Inductive Category System. The scheme depicts the inductively built category system used to classify statements (3). There are the main categories in the center; the first-order and second-order subcategories are indicated by decreasing color intensity. The figure does not include residuals, to which statements are assigned that could not be transferred to the existing categories.

To verify the consensus classification, we aim to categorize the qualitative data with a LLM. For this task, we chose the chat module of GPT 4.0 as a means to quantify the descriptive data. We described the category system and instructed ChatGPT to classify the collected data according to the provided category system. The entire procedure is text-based, and our starting point was a detailed description of the categories by Kaspar et al. (3), which was also the reference for the expert rater categorization. We introduced this description to ChatGPT and asked it to summarize the categories. We started to categorize on the main level with five categories. To finetune ChatGPT to interpret the statements as intended, we presented it with a few example statements. If classifications occurred that were incongruent with our understanding of the category system and the classification of the human raters, we would adjust the initially given instructions accordingly on what content belongs to which category and start a new chat. For instance, if the use of cardinal directions as a means to orient yourself was described and the statement is classified as “Space perception” by ChatGPT, but the category system marks the use of cardinal directions as belonging to the “Navigation” category. Then, we would change or expand the description of “Navigation” explicitly stating that using cardinal directions is part of “Navigation”. This way, we familiarized ChatGPT with our concept of the categories until the criterion of 10 classifications without a misclassification was met. Subsequently, we would classify the entire dataset and not interfere if misclassifications occur. The qualitative data was naturally separated by the week the statement was recorded. Yet, some statements are very comprehensive and include aspects that belong to different categories. Therefore, these statements have been segmented to assign different categories to the individual snippets. To still be able to differentiate between the categories, the phrases could not be assigned to multiple categories. Each was assigned to exactly one category. We limited the classification to the main level, which contains the categories “Space perception”, “Navigation”, “Belt experience”, “Feelings” and “Residuals”. To obtain an inter-rater reliability we compute the number of correctly classified statements divided by the total number of statements. Hence, by providing text-based explanations of the existing category system to ChatGPT, we can classify qualitative data in a way that enables quantitative analysis of collected statements.

## Results

First, we will report the average wearing and training times, and participants’ motivation to use the belt throughout the six weeks of training. The average wearing duration of the participants amounts to 9.205 hours per day (h/d; *SD* = 2.760 h/d). The active training duration was *M* = 2.270 h/d (*SD* = 1.738 h/d). Compared to a previous study (3), this represents an increase of approximately 42 minutes/day and exceeds the targeted 1.5 h/d. The reported wearing duration fluctuated between 8.714 h/d and 9.642 h/d throughout the training period without showing a clear trend. Motivation to wear the belt was highest in week 1, with an average of *M* = 4.295 (*SD* = 0.723) where 5 would indicate maximum motivation. It decreased constantly until week 5, *M* = 3.369 (*SD* = 1.182). After week 6, we observed a small incline (see Fig. 2). The average motivation during the training was *M* = 3.704 (*SD* = 0.984), indicating that most participants were motivated throughout the six weeks. Hence, the reported training duration and motivation indicate our participants’ high incentive to participate productively.

**Fig. 2.**
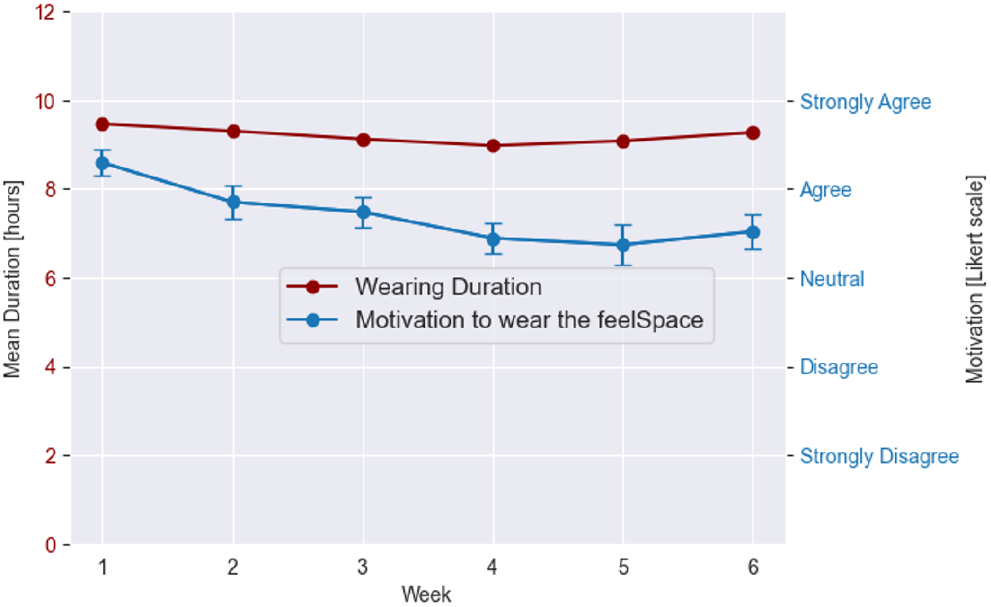
Belt Wearing Duration and Motivation. This figure shows how much participants wore the feelSpace belt and how their motivation developed over six weeks of training. The average wearing duration per week is displayed in red (left scale). The subjectively reported motivation to wear the belt is displayed in blue (right scale). The error bars of the blue line show the standard error of the mean.

### Aggregated Reports and Data

With the provided training parameters in mind, here we introduce the data collected throughout the training to illustrate a chronology of changes. We establish this chronology by classifying statements based on their content, allowing us to quantitatively identify trends in the data. In the following section, qualitative and quantitative data are presented together. We assume that categories reported more often in one week than another indicate a change in the respective domain. Therefore, we present clusters of categories and group various observations by their frequency over the training period. Similarly, if the rate of change of a numeric variable is high, this indicates that a substantial number of participants is experiencing a change that characterizes the present phase. Thus, we consider different training phases in sequence and base this order on the report frequency of categories and the rate of change in the numeric data.

### The Early Stage: Activation

In this section, we investigate the thematic distribution of statements in the first two weeks of the experiment. Statements assigned to the categories “Use of belt results in mental exhaustion”, “Wearing a correctly functioning belt causes irritations”, “More awareness for the environment”, “Belt indicates cardinal directions”, “Spatial relations between objects, locations, and streets”, “Ego perspective”, and “Mismatch” are most often early in the training phase and are grouped accordingly (see Fig. 3). In the early stage, there is an increased referencing of cardinal directions. Additionally, this stage is marked by active processing of the belt signal, heightened awareness of external information, and symptoms of mental effort. The participants appear constantly aware of the novel stimulus and their environment. Moreover, they are actively trying to incorporate it. In weeks 1 and 2, respectively, participants report “I’m more aware of the cardinal directions”, and “It [the feelSpace belt] is helping me to remember the cardinal directions in familiar areas”. Supporting this, the numeric data for conscious perception of cardinal directions is highest in week 1, *M* = 4.778 (*SD* = 0.641), then slightly decreases over the following two weeks. Besides an increase in their awareness of cardinal directions, participants also report relating objects in the environment to each other. Representative of this change in space perception is the following statement from week 2: “It’s easier to get the details right, especially with how the streets in Osnabrück are arranged”. We report that the numeric data estimating locations to each other increases by more than .5 Likert scale units from weeks 1 to 3. A Wilcoxon signed-rank test comparing the first and the last week further revealed a significant increase in estimating locations, *p* < .01 (see Table 1). The data suggest a more alert perception of the environment and the relations between objects in the environment. Moreover, our qualitative data demonstrates increased mental exhaustion and irritation caused by the correctly functioning belt in the first two weeks. One participant reports in week 1, “It [the feelSpace belt] is fairly exhausting to wear, also something of a change in habits […]”. The numeric data shows that the cognitive effort/concentration is the highest in week 2, *M* = 4.074 (*SD* = 0.675). It then decreases up to week 4 from “agree” to “slightly agree”, *M* = 3.519 (*SD* = 1.051). The trend suggests that signal processing becomes more effortless. Hence, in the early stage, participants can perceive mental conflicts caused by competing established representations and the augmented signal. Interpreting the augmented signal requires active processing during this stage, which may cause exhaustion. Reports on mismatches are very present in the first weeks, in week 3 a participant reports: “The hard cardinal information doesn’t align well with my personal map”. As a result of the augmented signal, participants compare the information to their existing knowledge, which may lead to competing representations. Interestingly, a few participants reported increased use of egocentric reference frames, while others reported that the belt information increases their awareness of cardinal directions and the environment. Moreover, multiple participants reported that they attempted to relate the objects and locations in their environment to each other more frequently. A Wilcoxon signed-rank test revealed that the numeric item regarding the estimation of the spatial relation of places towards each other increases significantly from weeks 1 to 6, *p* < .01 (see Table 1). Finally, the integration of the augmented input causes exhaustion and irritation. Due to the increased activity and the discovery of novel relations between objects in the environment, this stage is called the “Activation Stage”.

**Table 1.** Test Results of the Numeric Data. The table depicts in the first column the variables that were assessed, please refer to the data repository (see Data Availability) for the exact wording of the questions. The second and third columns depict the mean and standard deviation for the first week’s item assessment. The fourth and fifth columns depict the respective values for the sixth week. The test results column contains the test statistics (Stat) for the Wilcoxon signed-rank test and the corresponding *p*-value.

| Item | Week 1 |  | Week 6 |  | Test results |  |
| --- | --- | --- | --- | --- | --- | --- |
|  | M | SD | M | SD | Stat | p |
| How much did the belt restrict you. | 1.63 | .63 | 2.15 | 1.06 | 4 | <.01 |
| I was motivated to wear the belt. | 4.3 | .72 | 3.52 | 0.98 | 23 | <.01 |
| I am consciously perceiving the belt. | 3.04 | 1.06 | 2.63 | .88 | 45.5 | .13 |
| I perceive the transmitted information as a vibration. | 4.37 | .56 | 3.96 | 1.06 | 33 | .06 |
| I do not perceive the transmitted information as a vibration, but differently. | 1.74 | .9 | 2.44 | 1.25 | 48 | <.05 |
| After taking off the belt, I still perceive a sustained feeling of vibration. | 3 | 1.21 | 2.04 | 1.29 | 43.5 | <.05 |
| While wearing the belt, I am always aware of my location relative to my home. | 3.41 | 1.12 | 4.26 | .71 | 21 | <.01 |
| I am consciously concentrating on the belt, to use provided information. | 3.74 | .94 | 3.56 | .93 | 25.5 | .28 |
| It is easier with the belt than without the belt to orient myself in an unknown area | 3.67 | 1.24 | 4 | .88 | 40.5 | .25 |
| I find it easier with the belt to estimate the location of different places relative to other places. | 3.48 | 1.25 | 4.22 | .89 | 22 | <.01 |
| With the belt, I find it easier to estimate how streets are orientated relative to other streets. | 4.15 | .91 | 4.52 | .58 | 22.5 | <.05 |
| I possess a "cognitive spatial map" of my living environment. | 3.78 | 1.09 | 4.41 | .69 | 0 | <.01 |
| Since I started wearing the belt, I have perceived cardinal directions more consciously. | 4.78 | .64 | 4.63 | .74 | 5.5 | .28 |
| My spatial navigation abilities have improved subjectively since I started wearing the belt. | 2.85 | 1.03 | 4.07 | 1.04 | 14 | <.001 |
| When I take off the belt, my spatial navigation abilities get worse. | 2.15 | 1.17 | 3.22 | 1.19 | 16 | <.001 |
| When I am wearing the belt, my fellows react positively to it. | 3.52 | 1.19 | 3.7 | 1.23 | 49 | .29 |
| With the belt, I feel safer in an unknown area. | 2.63 | 1.24 | 3.44 | 1.31 | 15 | <.001 |

| Item | Week 1 |  | Week 6 |  | Test results |  |
| --- | --- | --- | --- | --- | --- | --- |
|  | M | SD | M | SD | Stat | p |
| How much did the belt restrict you. | 1.63 | .63 | 2.15 | 1.06 | 4 | <.01 |
| I was motivated to wear the belt. | 4.3 | .72 | 3.52 | 0.98 | 23 | <.01 |
| I am consciously perceiving the belt. | 3.04 | 1.06 | 2.63 | .88 | 45.5 | .13 |
| I perceive the transmitted information as a vibration. | 4.37 | .56 | 3.96 | 1.06 | 33 | .06 |
| I do not perceive the transmitted information as a vibration, but differently. | 1.74 | .9 | 2.44 | 1.25 | 48 | <.05 |
| After taking off the belt, I still perceive a sustained feeling of vibration. | 3 | 1.21 | 2.04 | 1.29 | 43.5 | <.05 |
| While wearing the belt, I am always aware of my location relative to my home. | 3.41 | 1.12 | 4.26 | .71 | 21 | <.01 |
| I am consciously concentrating on the belt, to use provided information. | 3.74 | .94 | 3.56 | .93 | 25.5 | .28 |
| It is easier with the belt than without the belt to orient myself in an unknown area. | 3.67 | 1.24 | 4 | .88 | 40.5 | .25 |
| I find it easier with the belt to estimate the location of different places relative to other places. | 3.48 | 1.25 | 4.22 | .89 | 22 | <.01 |
| With the belt, I find it easier to estimate how streets are orientated relative to other streets. | 4.15 | .91 | 4.52 | .58 | 22.5 | <.05 |
| I possess a "cognitive spatial map" of my living environment. | 3.78 | 1.09 | 4.41 | .69 | 0 | <.01 |
| Since I started wearing the belt, I have perceived cardinal directions more consciously. | 4.78 | .64 | 4.63 | .74 | 5.5 | .28 |
| My spatial navigation abilities have improved subjectively since I started wearing the belt. | 2.85 | 1.03 | 4.07 | 1.04 | 14 | <.001 |
| When I take off the belt, my spatial navigation abilities get worse. | 2.15 | 1.17 | 3.22 | 1.19 | 16 | <.001 |
| When I am wearing the belt, my fellows react positively to it. | 3.52 | 1.19 | 3.7 | 1.23 | 49 | .29 |
| With the belt, I feel safer in an unknown area. | 2.63 | 1.24 | 3.44 | 1.31 | 15 | <.001 |

**Fig. 3.**
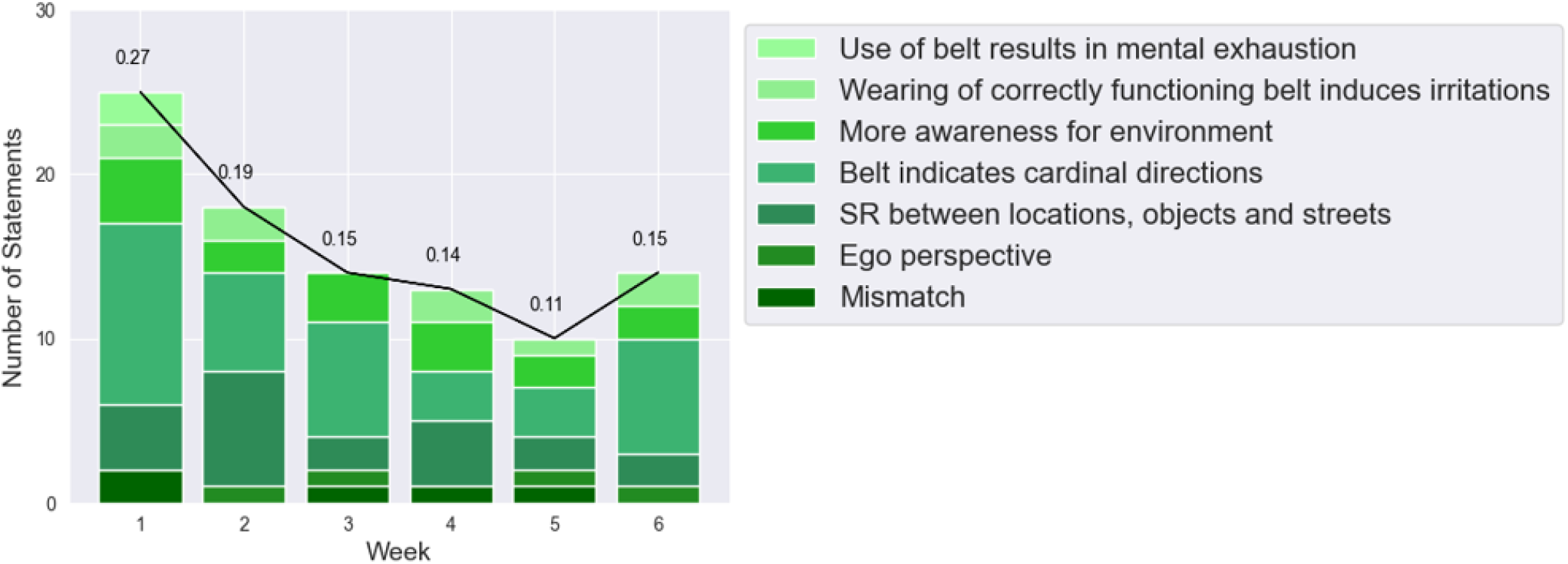
The Activation Stage. The abscissa corresponds to the weeks of training and the ordinate to the number of statements collected each week. The color coding of the bars separates the different categories, each color corresponds to one category, as indicated by the legend. The decimals above the bars represent the proportion of statements collected throughout the respective week regarding the total number of statements.

### The Intermediate Stage: Knowledge Acquisition

In the following, we continue linking quantitative and qualitative data to analyze perceptual changes after the early stage. Reports on corrections regarding the mental representations of our participants are increasingly more present from week 3 onward. Specifically, categories that contain most statements during the intermediate section of the training are “Mental map is used for navigation”, “More detailed representation of space”, “Enlarged mental map”, “Relation of locations and objects to oneself” and “Correction” (see Fig. 4). For example, one participant reports in week 4: “My mental cognitive map is not as warped as last time, or maybe I got better at adjusting it on the fly based on the cardinal direction information”. The participants replace inaccurate information in their mental map. Thereby, participants’ mental representation of space is changing. This development is indicated by statements involving descriptions of a more detailed representation of space, an enlarged mental map, and relating the location of objects to themselves. As such, the participants report: “I have a clearer picture of my location in my mental map than before” indicating a more detailed representation of space, or “I always move at a certain angle relative to the street where I live” as an instance of how participants relate objects to their position. The findings of implementations happening on mental representations align with the developments of the numeric data. The number of statements referring to the orientation of streets towards other streets increases in particular after week 3. Yet, the results of a Wilcoxon signed-rank test revealed a significant difference between weeks 1 and 6, *p* < .05, suggesting the development continues in the last third of the training (see Table 1). Further, we find that the awareness of the participants’ position concerning their home has increased by more than .5 on the Likert scale from weeks 2 to 3. The estimation of street alignments by .3. Moreover, participants learn to interpret new patterns while navigating. Overall, the intermediate stage is marked by increasing reports on implementations of the mental representation of space and less mental exhaustion or irritation. The participants learn to integrate new knowledge into existing representations and start acting upon it more successfully, thereby earning its name, “Knowledge Acquisition Stage”.

**Fig. 4.**
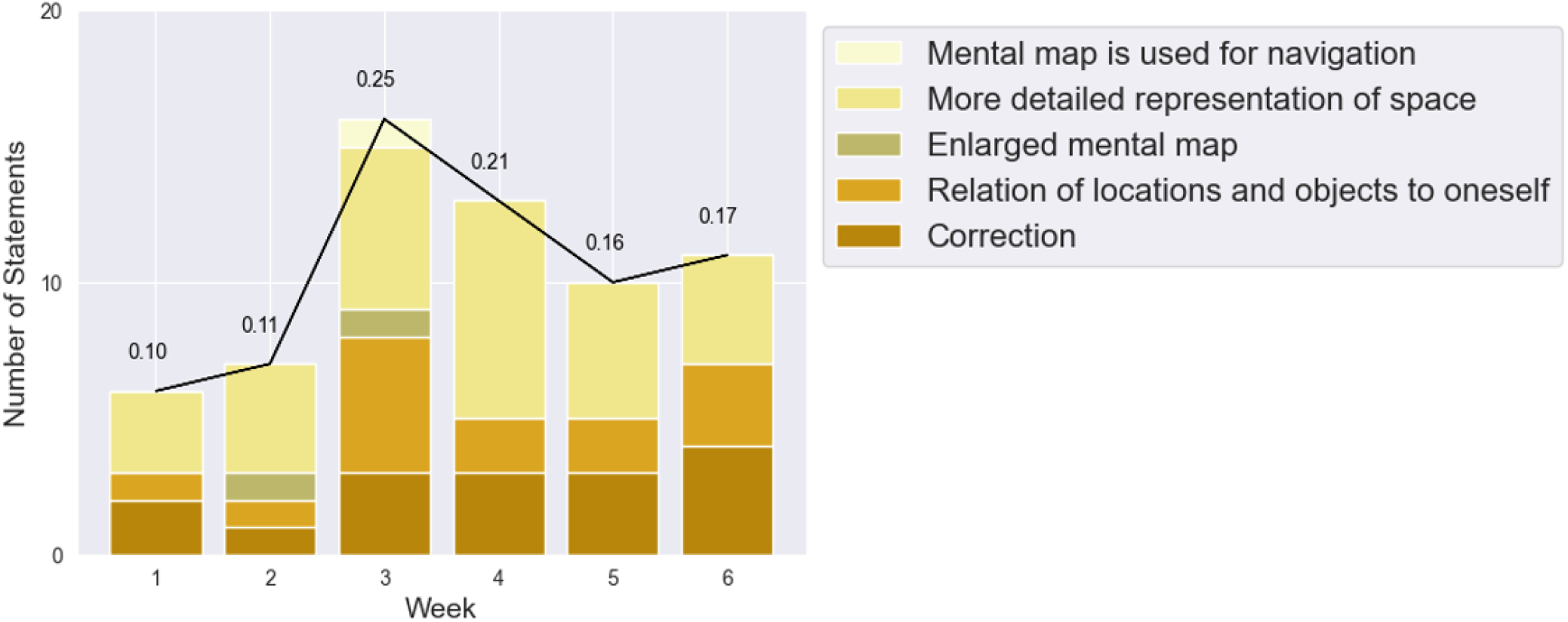
The Knowledge Acquisition Stage. The abscissa corresponds to the weeks of training and the ordinate to the number of statements collected each week. The color coding of the bars separates the different categories, each color corresponds to one category, as indicated by the legend. The decimals above the bars represent the proportion of statements collected throughout the respective week regarding the total number of statements.

### The Late Stage: Deep Integration

Toward the end of the training, several participants reported that the belt signal substantially facilitates their spatial navigation; this change indicates entering the late stage. Categories that occur most often late in training are “Belt facilitates navigation”, “Aerial perspective”, “Something is missing without the belt”, “Feeling of insecurity without the belt”, “Space perception beyond visibility”, “Alignment/orientation as a property of objects”, and “Spontaneous navigation with little reflection” (see Fig. 5). Moreover, the alignment as a property of an object is in particular present in participants’ reports, yet this aspect arises already at the beginning of training before week 5. An instance of this category is the statement: “I have more awareness of the slight tilts in the angle of streets”. There is also an increase in the number of participants who report feeling insecure or incomplete without the belt. In weeks 4 and 6 two participants reported: “Yesterday, while I was not wearing the belt at work I was constantly feeling its absence”. Moreover, a participant reports: “I feel more confident in unfamiliar environments and try to find my way on my own without looking directly at maps”. Hence, they notice the emotional relevance of the belt for their navigation and orientation. Complementary to this observation, we see a significant increase in the numeric data. Two items, spatial navigation ability and the feeling of security in unknown environments increased by more than .8 from week 1 to 6. A Wilcoxon signed-rank test revealed that these increases were significant, *p* < .001 (see Table 1). Within the same time the estimated ability to navigate in unknown environments improved by .33, however, this increase was not significant, *p* = .25 (see Table 1). Taken together, these observations show that participants integrate the belt signal into their navigation strategies.

**Fig. 5.**
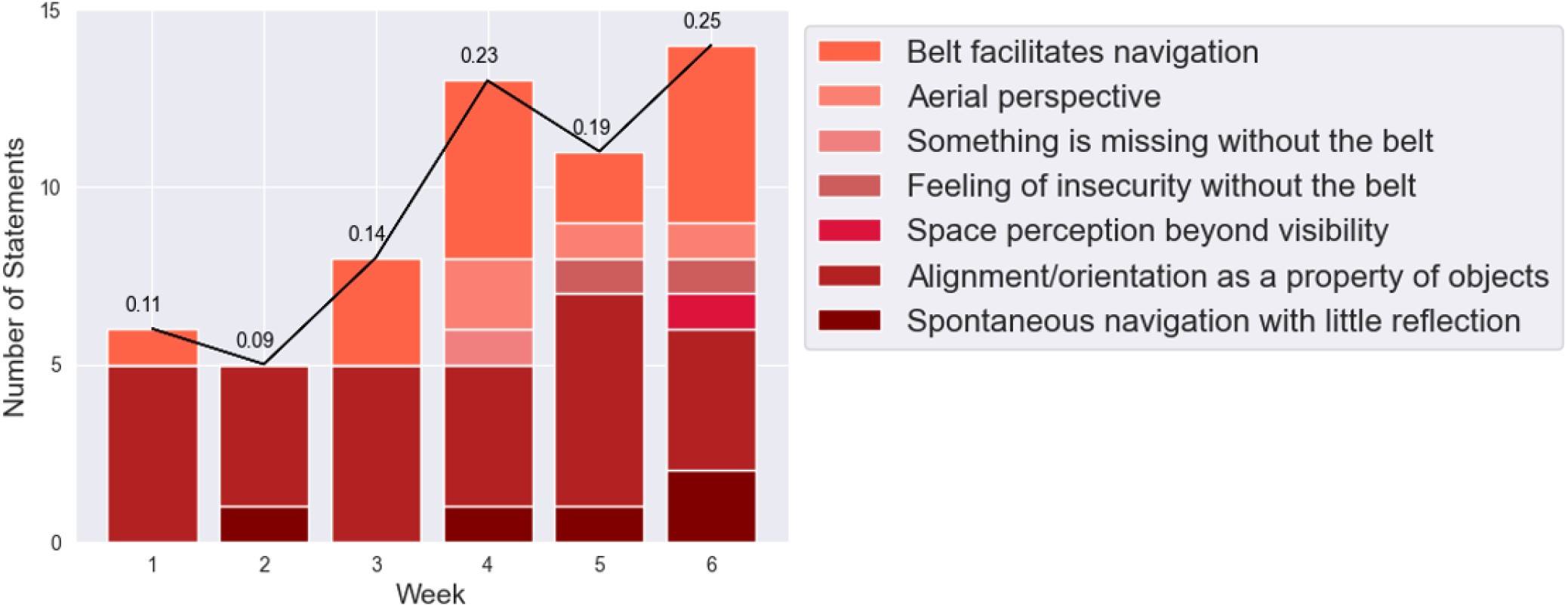
The Deep Integration Stage. The abscissa corresponds to the weeks of training and the ordinate to the number of statements collected each week. The color coding of each bar separates the different categories; each color corresponds to one category, as indicated by the legend. The decimals above each bar represent the proportion of statements collected throughout the respective week concerning the total number of statements.

In the last week, space perception beyond visibility occurred indicating a sense of space that is not dependent on vision. “Now I have the feeling that I perceive my surroundings even if I can’t see them directly, e.g. I know the directions of the cars at the intersection without thinking about it, and without looking out of the window”. The Paticipant’s statement suggests that augmented knowledge becomes an integrative part of perception. In the second half of training, a few statements come up about an aerial perspective on space (see Figure 5), and in week 3, one person reports using their mental map for navigation, stating: “I can call up the “mental map” more often and more easily, or it comes automatically more often”, thus, we witnessed attempts of deploying globally-oriented navigation. From week 3 onward we see an increase in reports describing the belt as facilitating navigation. In week 5, a participant stated: “I’m using the information the belt provides consciously to navigate especially through unknown neighborhoods”. The participant mentions that the incorporation of the signal happens actively. More automatic usage of the signal is entailed in the category “Spontaneous navigation with little reflection”, instances are the following statement:

“Orienting myself in the world became easier. Finding my way from A to B has become a bit more automatic and less dependent on planning ahead” or “Basically just using it [the feelSpace belt] as a signal without too much thought or effort behind it”. The automatic navigation is reported less than the category “Belt facilitates navigation”. However, both categories start occurring approximately simultaneously, and the reports increase towards the last third of the experiment. The Likert scale data assessing the conscious belt perception is aligned with such qualitative observations, the highest rating is *M* = 3.110 (*SD* = 1.085) in week 2. Afterward, we see a decrease until week 6 with *M* = 2.630 (*SD* = 0.884). Finally, in the first week, we asked our participants whether they possessed a cognitive spatial map of their environment, which 21/27 confirmed and 6/27 denied, in the last week 23/27 confirmed. The item in the questionnaire assessing the participants’ possession of a cognitive map was growing continuously from week 1, *M* = 3.778 (*SD* = 1.086) until week 6, *M* = 4.407 (*SD* = .694). A Wilcoxon signed-rank test comparing the first and last week revealed that this increase was significant, *p* < .05 (see Table 1). Generally, participants feel more secure with the belt and estimate their navigation ability to be improved, they also emotionally rely on the feelSpace belt. These findings indicate that participants deploy the belt signal in navigation and deeply integrate the augmented information into their perceptual processes and spatial cognition.

### The activation-acquisition-integration model of sensory augmentation

The data presented above indicate elaborate dynamics during the training period. At all times, statements of any category might occur. Nevertheless, a systematic shift in dominating statements is observable. Early in the training phase statements emphasize unspecific effects like more awareness, mismatch, and exhaustion. Then, the focus shifts to the knowledge supplied by the sensory augmentation device. Only late in the training is a deeper integration and emotional evaluation more prominent. This shift in focus in participants’ reports is best captured by a three-stage model: activation, acquisition, and integration (see Fig. 6).

**Fig. 6.**
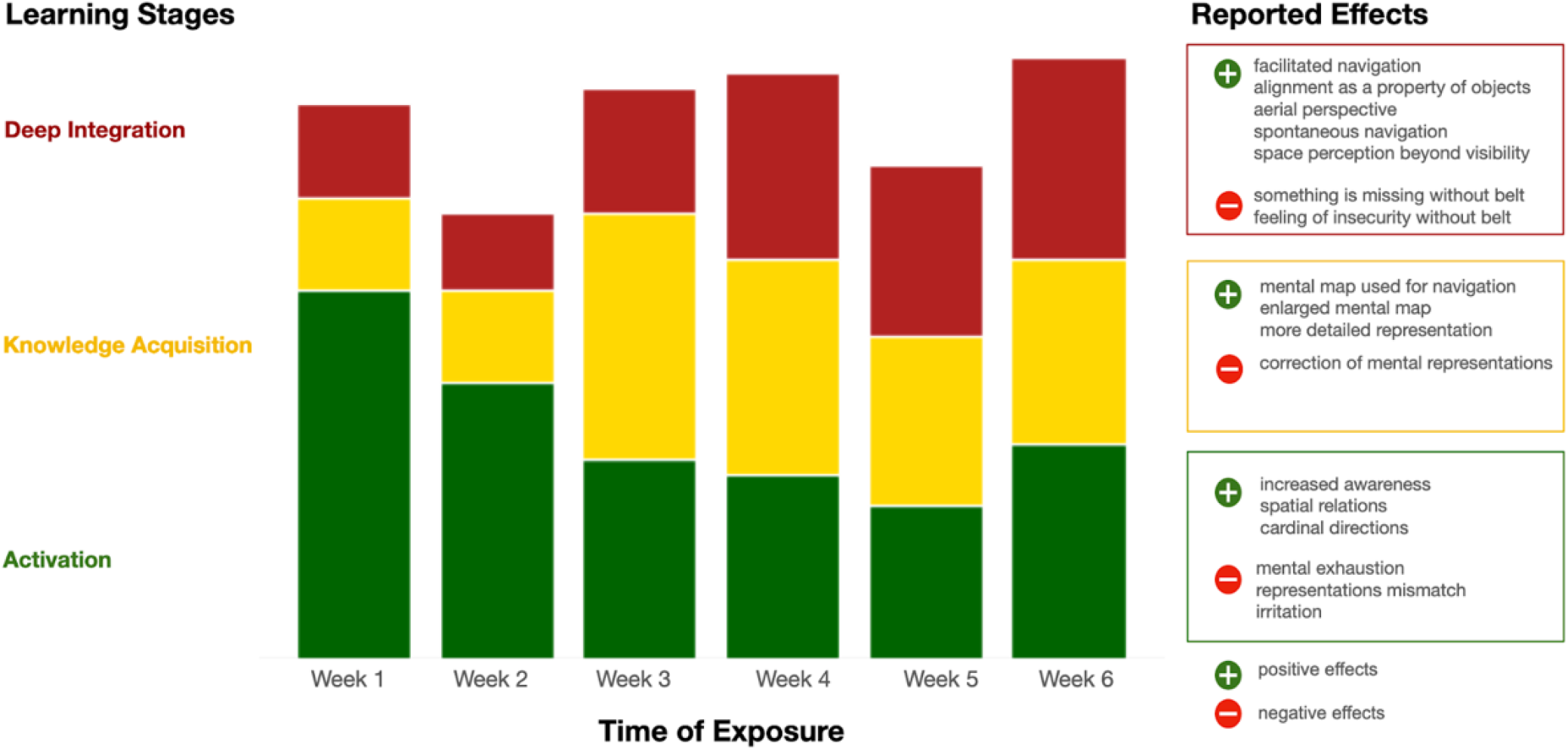
The Activation-Acquisition-Integration Model of Sensory Augmentation. The scheme illustrates the model presented in the previous section, consisting of the activation, knowledge acquisition, and deep integration stages. The abscissa indicates the time of exposure to the augmented input and the ordinate shows the stages on the right-hand side and the corresponding changes on the left-hand side. The focus of reports shifts systematically during the training, creating three stages.

### Classification Verification using ChatGPT

To address the problem of objectively classifying responses to open questions, we employ a widely available language processing and generation tool: ChatGPT. Our rationale was to investigate whether the representation of the category content can be conveyed to a LLM. Specifically, we wanted to know whether a LLM would reproduce the human rater classification. The ChatGPT classification was performed with 475 segments; the difference in the number of statements (compared to the total number of statements of 522) is because the ChatGPT classification was performed on the main category level, as indicated in the Methods section. The segmentation of statements is more fine-grained with increasing specificity of the categories. Hence, there are more samples on the second-order category level than on the main category level. The distribution of the interrater and ChatGPT classification is presented in Fig. 7. The resulting distribution of the categories “Navigation”, “Belt experience”, and “Residuals” match up to ± 0.6%, and the “Space perception” category is slightly larger. The “Feelings” category is underrepresented in the ChatGPT classification and loses almost half its count, which amounts to 2.6% of all statements. The total count of the main level category assignments of consensus and Chat-GPT classification are largely aligned.

**Fig. 7.**
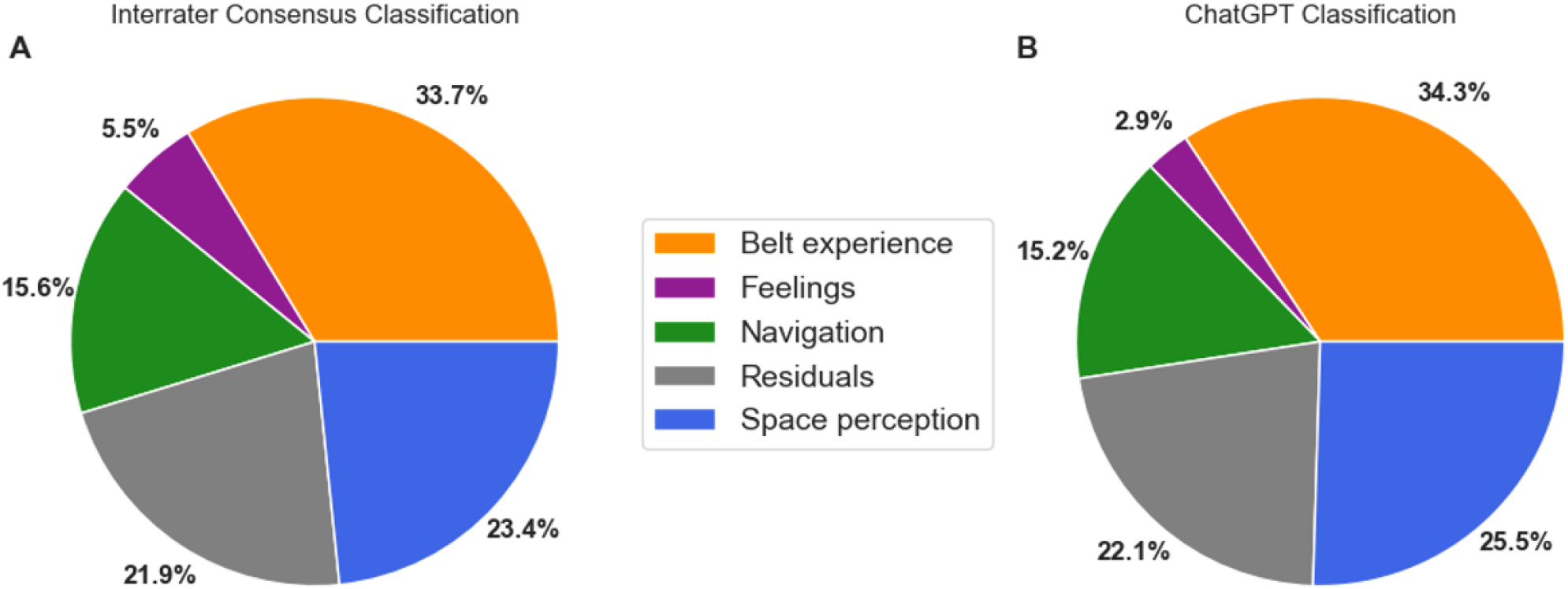
Category Distributions of Qualitative Data. This plot shows the count of aggregated classification of participants’ responses into main categories. Panel A shows the expert rater consensus classification, and panel B shows the same data but classified by ChatGPT. The percentages indicate the proportions of the respective categories relative to the total number of statements.

As a next step, we compare the congruence of single statement classification. ChatGPT classified 84.41% of the statements matching the interrater consensus classification. Without the Residuals category, which mostly includes brief statements on topics unrelated to our research questions, the accuracy of ChatGPT classification is 81.67%. Although it is a rather high value, it indicates that the excellent match in the overall distribution of categories reported in Figure 7 hides some disagreement. An analysis by a confusion matrix uncovers that most confusion arises between the “Space perception” and “Navigation” categories (see Fig. 8). Seventeen out of 74 statements from “Navigation” were classified as “Space perception” and 20 out of 101 statements from “Space perception” were classified as “Navigation”. The statements that have been assigned to the “Feelings” category were often misclassified as “Belt experience”. The ChatGPT model has reproduced the distribution of categories predominantly accurately, i.e. misclassifications occur at a moderate rate only, and the direction of confusion is symmetric. However, concerning single statements, the discriminatory power across categories could be improved. Overall, we conclude that the ChatGPT classification employed in our approach is satisfactorily congruent to the human rater classification and allows a classification that is not dependent on specific individuals.

**Fig. 8.**
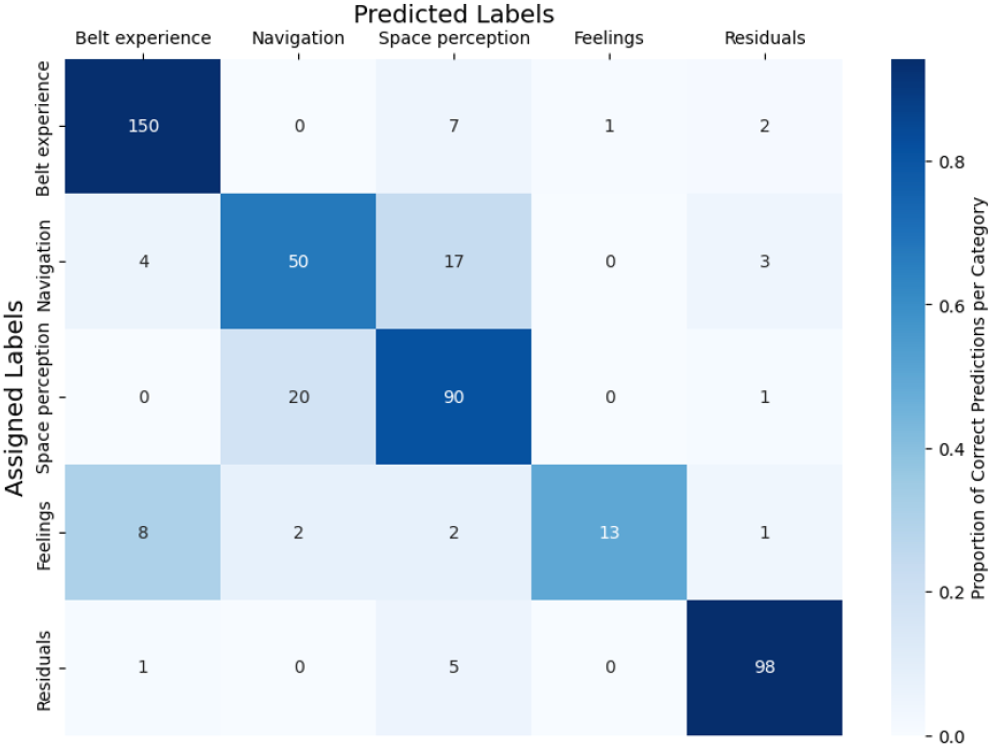
Figure 8. Confusion Matrix of ChatGPT Classification. The confusion matrix represents the congruence between the assigned categories by human raters and the predicted categories by ChatGPT. The color indicates how many statements have been correctly classified per category. The digits in each cell correspond to the number of statements in the respective category.

## Discussion

In this study, we investigate the temporal dynamics of sensory augmentation, namely how the augmented information of the feelSpace belt affects space perception and navigation during training with the device. Our results have led us to identify three stages that describe the changes that occur during exposure to an augmented sensory input. As a result, we propose the activation-acquisition-integration model (see Fig. 6), which describes which perceptual and action-related changes participants reported to occur predominantly at which point in time. Moreover, the results show that most participants follow a temporal development from an effort-intensive activation stage over a knowledge acquisition phase toward a more integrated and automated application of the knowledge gained through the augmented sense. The framework was developed and tested with the feelSpace belt, however, we speculate that the temporal dynamics are not specific to this device but describe changes related to sensory augmentation in general. Hence, the likely initial effects include exhaustion from actively processing the augmented information and heightened sensory perception awareness. Over time, the application becomes less error-prone and requires fewer resources as understanding the signal becomes more intuitive. In conclusion, the temporal dynamics of learning an augmented sense proceed in three stages with distinctive characteristics in the degree of effort required for processing and successful application exhibited.

### The Assessment of Qualitative Data

When investigating perceptual changes, researchers have to face numerous well-established problems. Here, we did not physically measure behavioral changes but estimated perceptual changes based on questionnaire data and interviews. Thus, we encounter subjective biases that come along with self-reported data. Such data is prone to individual differences in self-perception that lead to measurement errors (16, 17). We face this problem by adopting a larger sample size. At the same time, we can refer to previous studies that align with our findings of perceptual changes and emotional states induced by the feelSpace belt (2, 3, 21). Complementary to the self-reported data, also Schmidt and colleagues (11), with behavioral experiments and König et al. (14) assessing neurophysiological activity, confirm the effects of wearing the feelSpace belt. Thus, while our investigation relies on self-reported data, results have been consistent throughout different studies with diverse methodologies.

When analyzing qualitative data, classifying statements based on their content poses a risk for subjective biases. The category system for classification had been introduced and tested previously, which makes it an established approach (3). To evaluate the discriminatory power and accuracy of the human raters, we implemented a classifier based on Chat-GPT. By describing the categories to ChatGPT and creating a statistical model of the categories, we deploy an approach to autonomously classify the data and take the obtained distribution of categories as a reference for the accuracy of the human-based categorization. The congruence between Chat-GPT and human rater classification is high but leaves room for improvement. We suggest including more policies on treating specific types of statements to improve the classification accuracy further. The statement “same as previous week” should be classified as the statement of the previous week. However, this statement has been assigned to the “Residuals” category using the current approach. Such formal adjustments would increase the precision of the ChatGPT classification. Additionally, the category system could be adjusted, increasing the discriminatory power between the category “Space perception” and “Navigation”. For now, the category similarity between “Belt indicates cardinal directions” which is part of “Navigation” and other categories that belong to “Space perception” like “More detailed representation of space” lead to some confusion. It is questionable whether the category “Belt indicates cardinal directions” belongs to “Navigation”, as perceiving cardinal directions through the belt does not necessarily involve the application of the signal, which arguably is necessary for navigation. Hence, a refinement of the subcategories of the system reflects a possibility to decrease the number of misclassifications.

The autonomous classification approach itself exhibits pitfalls too. A subjective element in the verification approach is represented by the active alignment of the representation of ChatGPT with the raters’ understanding of the category system. As such, we might transfer certain biases to the ChatGPT classification. Furthermore, the intransparent nature of LLMs impedes the detailed mechanistic understanding of the autonomous classification method. On the other hand, ChatGPT is a tool accessible to a large audience, which in principle, enables the verification of our results. Despite the named downsides, the classification approach with Chat-GPT improves the consistency of the category assignments. It provides a qualitative data classification tool that can be applied in any lab to various demands concerning content analysis.

### Training Duration

In this study, we observed perceptual changes throughout six weeks of training with the feelSpace belt. Yet, some changes like “Space perception beyond visibility” occur rather late in the 6-week training period, suggesting that prolonged training would exhibit further development. Previous studies on the feelSpace belt investigated effects for six (2, 21) and seven weeks (3) with comparable results. Nevertheless, experiments with substantially extended training periods appear desirable. Moreover, Levy-Tzedek and colleagues (22) emphasized that including younger participants might be desirable as it has been shown that children are better at learning an augmented sense. Therefore, to study lasting and intuitive handling of augmented senses, we suggest that including younger age groups and a substantially longer training period are promising.

Early studies of sensory substitution started with rather short training. For instance, the pioneering study trained participants for up to 40 hours with a vision-to-tactile substitu-tion device that projects shape information to the back of blind participants (7). After only 10 hours of training, participants correctly identified an object drawn from a pool of 25 different shapes. In a follow-up study by Kaczmarek et al. (23), sighted blindfolded participants achieved recognition of 79.8% across four shapes after one training session with a tongue stimulation unit. Jones and colleagues (24) and Lozano and colleagues (25) investigated how quickly participants adapted to tactile stimulation on the waist by vibrating motors and electro-tactile stimulation on the tongue, respectively, in both cases, a few hours of training allowed the participants to identify simple patterns from the tactile stimulation. We can, therefore conclude that people can interpret simple patterns from tactile augmented input relatively quickly. Longer observation periods took place in the studies by Amedi and colleagues (10) and Badke et al. (26). Amedi et al. (10) trained sighted participants for 20 days with the “vOICe”, a device that translates visual shapes into an auditory signal. After that time, the subjects were able to map sounds to features and identify objects on an “expert” level. Test subjects in Badke et al. (26) trained for 1 week under clinical conditions and another 7 weeks as a home exercise. The subjects had balance deficits and were supported by a device that gives vibrotactile balance support on the thigh. After the training, the participants reported improved balance, gait function, balance confidence, and quality of life. A recent study (27) investigated the effect of vibrotactile balance support throughout a training period of 8 weeks with a session lasting one hour each day. The balance of the subjects improved significantly, and even 6 months post-training, an improved performance was detectable. They also recorded fMRI before and one week after training to assess neurophysiological changes. They concluded that the brain activity during vestibular stimulation moved from cortical regions pre-training to brainstem and cerebellar regions posttraining. This suggests a more automatic control after the training. Moreover, the complexity of the stimulus and the task greatly influence the time required to incorporate the stimulus into the performance. Thus, a higher stimulus or task complexity will require longer training periods. Altogether, these findings show that sensory substitution devices lead to quick adaptation and integration of the provided information and, when applied for longer periods, may result in lasting changes beyond the training period.

A further factor that might influence training time and performance is presented in the cognitive and physical condition of people using sensory augmentation devices. The EyeCane (13, 28) is a sensory substitution device that gives auditory feedback on the visual modality. It was tested on blind and blindfolded sighted subjects concerning distance estimation, navigation, and obstacle avoidance. After only 3-5 min of introducing the device, the participants mastered the given tasks. Further, Chebat et al. (29) equipped sighted, blindfolded, and blind participants with a vision-to-electrotactile tongue stimulation device and asked about the properties of an object (distance: near, far; size: small, large), all subjects were able to classify the objects correctly. In both studies, the visually impaired subjects outperform the (sighted) blindfolded subjects. Maidenbaum and colleagues (28) and Chebat et al. (29) reported effective tactile substitution on navigation-related domains after a brief introduction period to the respective devices. In the current study, we also observe an initial sense-making of the augmented input, which is then processed and integrated into actions rather quickly. However, the 3-stage model of the underlying temporal dynamics shows that established navigation strategies may be challenged due to having the augmented signal at their disposal. Besides the contradictory information from the feelSpace belt and the established navigation strategies, the variety of demands in daily spatial orientation has led to a longer adaptation period in the present study. The stronger performance of blinded subjects could also be attributed to a stronger focus on the tactile domain in the life of blind subjects (30) in the named studies (13, 28, 29). However, the absence of some sensory information may also simplify the adaptation to the tactile translation of spatial information (31, 32). Thus, overlapping modalities for similar pieces of information may lead to more conflicts, resulting in increased time required to adapt to an augmented input and an initially decreased performance.

### Sensorimotor Theory of Perceptual Consciousness

Embedded in the framework of sensorimotor contingencies (1), we investigate a novel sense provided through the feelSpace belt. As the feelSpace belt indicates the position of the magnetic north, it can provide humans with information that is otherwise not directly accessible (33, 34). By learning to use a novel sensory signal, such as provided by the feelSpace belt, the input is integrated into the perceptual experience. Our research shows how the learning process manifests and how perception and action are affected until, eventually, SMCs are formed and manifested.

The acquisition of SMCs is a well-established approach to explaining conscious perceptual experience by law-like patterns that connect action and perception. However, the actual process of learning such SMCs was undocumented. Therefore, we attempt to fill this gap by introducing the hierarchical stages we observed during the training period. Here, we examine whether our findings align with the essential nature of SMCs and their acquisition through a reciprocal relation of action and sensory feedback (35). At first, during the training, the functionality of the augmented domain diminished, and the learning was effort-intensive. Yet throughout the entire training period, we collected evidence that participants either actively or passively compared the directions indicated by the belt and their intuition for where the magnetic north or other locations are located. We observed a tendency from active to more passive comparison from start to end of training. In week 3, a participant states: “[…] my cognitive map is warped and imprecise and the hard cardinal information doesn’t align well with my personal map […]” and in week 6 “[…] being aware of the signal and maybe subconsciously comparing it to previous experiences[…]”. Another statement providing interesting insights into the learning process is “Sometimes when I go around a bend, I think for a moment that the signal has not changed. But when I think about it, I realize that the position of the vibration on my belly has changed but the direction of the signal has not”. The statement might indicate a conflict between automatic processing and active monitoring of expected sensory feedback. This is congruent with our data on cognitive effort and conscious processing of the belt signal. Within the sensorimotor theory, our findings confirm the prediction of repeated probing of the environment through sensory feedback.

During the early learning stage, participants attempt to apply the novel information despite experiencing mismatches and possessing incomplete information. For example, one participant reported after the first week: “It’s easier for me to orient myself in a new environment and to take new paths. In addition, I can better remember routes in new cities, and especially return routes are easier to remember”. The participant applied the provided information at least partially successfully. At other times, participants appear to struggle: “With the additional information the belt is giving me, I feel like the two systems of navigation are not yet integrated (week 1)”. With the continued probing of the signal, the participant’s navigation changed several times when considering the development of single participants, they described how problems they experienced while processing the feelSpace belt signal gradually improved over time. As such, the following week, the author of the previous statement wrote: “The belt does no longer disrupt my previous navigation technique but kind of co-exists”, eventually in week 6 “Before I only used my starting point or landmarks as orientation, but with the belt, I started taking cardinal directions into account and developed some kind of spatial map”. We observed that participants were probing their incomplete knowledge while paying attention to sensory feedback. Participants constantly monitored the incoming feedback, improving their success at responding to novel information and, eventually, automating the perception-response process. The feedback loop closely resembles the framework of SMCs (1; see Introduction), and seems to rely on proactive learning to become automated.

### Future Directions

To increase the temporal precision of the predictions that can be derived from the introduced paradigm, the collected data should be leveraged by self-reports. This is a far-reaching goal, as identifying neural activity or changes relating to specific qualia has not been accomplished yet (36). Analyzing further why there are clear interindividual differences in the acquisition and velocity of learning the feelSpace-related SMCs could contribute to refining the Activation-Acquisition-Integration Model. Moreover, temporal dynamics should be observed over a longer training period because the development within the deep integration stage was only visible after 5-6 weeks of training. With continued training further developments are likely to appear regarding domains that are not directly connected to the augmented sense, for instance, the mental representation of a city or the way one would coordinate locomotion during an exercise. Overall, future research can employ the activation-acquisition-integration model and its underlying methodology to make self-report data more accessible and interpretable for analysis, especially when mixed data types are available.

### Conclusion

In this study, we formulated a framework for the temporal dynamics of learning to integrate sensory augmented input into action and perception. As proposed by O’Regan and Noë (1), we found that the participants report perceptual changes that correspond to the acquisition of tactile feed-back on cardinal directions induced by the training with the feelSpace belt. The learning process was first characterized by resource-intensive processing. Then, the new signal was actively integrated together with existing senses and potential mismatches. Later, the signal became increasingly more intuitive and efficient to process. The resulting Activation-Acquisition-Integration model aims to summarize these findings, thereby providing a framework to compare and further investigate the dynamics of augmented learning. Further, we addressed the problem of inadequate physical measuring procedures for recording perceptual changes by analyzing self-reported data with a category system and ChatGPT as a means of content analysis.

## Author Contributions

Conceptualization, V.S., and P.K.; methodology, V.S., and P.K.; software, V.S., validation, V.S., P.K., and S.M.; formal analysis, S.M.; investigation, V.S.; resources, P.K.; data curation, V.S., and S.M.; writing—original draft preparation, S.M., and V.S.; writing—review and editing, V.S., S.M. and P.K.; visualization, S.M.; supervision, P.K., and V.S.; project administration, P.K.; funding acquisition, P.K. All authors have read and agreed to the published version of the manuscript.

## Review Board and Consent Statement

The study was conducted in accordance with the Declaration of Helsinki and approved by the Ethics Committee of Osnabrück University (protocol code: 4/71043.5, date of approval: 5 January 2021). Informed consent was obtained from all subjects involved in the study.

## Data Availability

All data and analysis scripts are available at [OSF](https://osf.io/n3qsc/). Additionally, the repository contains transcripts of the daily and weekly questionnaires and the transcript of the ChatGPT classification.

## ACKNOWLEDGEMENTS

We thank Rabia Dilawar, who contributed to the human rater data classification and helped us establish useful guidelines for edge-case scenarios. Similarly, we want to thank Dr. Sabine U. König for providing us with support regarding crucial materials and underlying concepts or theories.

